# Investigation of the *in vitro* and *in vivo* activity of the type IV pilus extension ATPase BfpD

**DOI:** 10.1101/2020.05.28.120931

**Authors:** Jinlei Zhao, Shahista Nisa, Michael S. Donnenberg

## Abstract

Type IV pili (T4Ps) are multifunctional protein fibers found in many bacteria and archaea. All T4P systems have an extension ATPase, which provides the energy required to push structural subunits out of the membrane. We previously reported that the BfpD T4P ATPase from enteropathogenic *E. coli* (EPEC) has the expected hexameric structure and ATPase activity, the latter enhanced by the presence of the N-terminal cytoplasmic domains of its partner proteins BfpC and BfpE. In this study, we further investigated the kinetics of the BfpD ATPase. Despite high purity of the proteins, the reported enhanced ATPase activity was found to be from (an) ATPase(s) contaminating the N-BfpC preparation. Furthermore, although two mutations in highly conserved *bfpD* sites led to loss of function *in vivo*, the purified mutant proteins retained some ATPase activity, albeit less than the wild-type protein. Therefore, the observed ATPase activity of BfpD was also affected by (a) contaminating ATPase(s). Expression of the mutant *bfpD* alleles did not interfere with BfpD function in bacteria that also expressed wild-type BfpD. However, a similar mutation of *bfpF*, which encodes the retraction ATPase, blocked the function of wild-type BfpF when both were present. These results highlight similarities and differences in function and activity of T4P extension and retraction ATPases in EPEC.

## Introduction

Type IV pili (T4Ps) are the most prevalent type of fimbria, found throughout gram-negative bacterial phylogeny, in some gram-positive bacteria, and in archaea. These pili perform a wide variety of functions including surface adherence, aggregation, twitching motility, DNA uptake, and electrical conductivity [1]. T4Ps are assembled by a complex machine that, in gram-negative bacteria, spans the inner and outer membrane [2]. T4P systems share similar assembly machinery and biogenesis mechanisms with Type 2 secretion systems (T2SSs) and archaeal flagella systems [3]. T4Ps are composed primarily of a major pilin protein, which, along with minor pilins, must be extricated from its inner membrane location as an integral transmembrane protein to polymerize into a helical extracellular fiber. An extension ATPase, present in all T4P and related systems, presumably converts chemical energy into mechanical energy, providing the work required for this task. For example, in *Vibrio cholera*, the T2SS ATPase EpsE is required for function of a putative pseudopilus, which is proposed to act as a piston during exoprotein translocation across the outer membrane [4]. An inner membrane component of this T2SS system, EpsL, was found to interact with EpsE and activate its ATPase activity 30-130-fold [5–6]. The *in vitro* ATPase activity of EpsE can also be increased by artificially enhancing its ability to form a hexamer, which is the natural state *in vivo* for T4P and T2SS ATPases [4].

Diarrhea is the second leading cause of death in children during their first five years, accounting for 9% of all childhood mortality in 2013 [7]. Particularly in Africa, infection with typical Enteropathogenic *Escherichia coli* (EPEC) strains is associated with a high risk of death [8]. The pathogenesis of typical EPEC infection involves a T4P called the bundle-forming pilus (BFP). Expression of BFPs leads to bacterial aggregation and to localized adherence (LA), where bacterial microcolonies formed by bundling of BFP adhere non-intimately to epithelial cells [9–10]. It is believed that typical EPEC strains use BFPs to colonize epithelial cells and more efficiently deliver effector proteins via their type 3 secretion systems therein [11].

We previously demonstrated that the cytoplasmic amino terminus of BfpC (N-BfpC), a structural homologue of EpsL and PilM of *Thermus thermophilus*, and the cytoplasmic N-terminus of BfpE (N-BfpE), homologous to the polytopic inner membrane protein present in all T4P and related systems, could be co-purified with hexahistidine-tagged BfpD [12]. Furthermore, the ATPase activity of BfpD was increased approximately 3-fold in the presence of both N-BfpC and N-BfpE, suggesting allosteric changes upon interactions among them. However, the above study did not investigate whether one or both proteins caused the stimulatory effect on ATPase activity, nor did it measure the classical enzymatic parameters V_max_ and K_m_. In this study, we sought to further the published investigations.

## Materials and methods

### BfpD, N-BfpC and N-BfpE expression and purification

All bacterial strains and plasmids used in this study are listed in S1Table. Full-length BfpD was overexpressed in *E. coli* BL21(DE3) as an N-terminal hexahistidine fusion protein. In short, the *bfpD* gene was amplified from plasmid BfpD-Hcp1. To make BfpD-Hcp1, codon-optimized *bfpD* was fused with Hcp1 [13] at the C-terminus and cloned into pET151/D-TOPO. In pBfpD-Hcp1, *bfpD* was also fused at the N-terminus with 6xHis - V5 epitope - TEV. Furthermore, the second residue was mutated from L to V to facilitate cloning. Primers BfpDNcoI and BfpDXhoI (S2Table) were used to amplify codon-optimized *bfpD*, which was cloned into the NcoI/XhoI sites of pET30a plasmid to make pJZM005. *E. coli* BL21(DE3) carrying plasmid pJZM005 was grown at 37°C in LB medium to an optical density at 600 nm (OD_600_) of 0.6 and induced with 1 mM isopropyl β-D-1-thiogalactopyranoside (IPTG) at 16°C overnight. Cells were harvested by centrifugation, lysed through sonication and purified by affinity chromatography on nickel resin. The resultant elution components were mixed and dialyzed against buffer [20 mM Tris–HCl, pH7.6, 100 mM NaCl]. When required to achieve further purification, a HiPrep 16/60 Sephacryl S-300 column was used.

Plasmid pRPA302 was used for the expression and purification of BfpC cytoplasmic amino terminal residues 1-164 (N-BfpC) as previously described [14], except that cells were lysed through sonication and a HiPrep 16/60 Sephacryl S-100 column was added for some preparations. Cytoplasmic amino terminal residues 1-114 of BfpE (N-BfpE) was cloned into pTEB68 as a NcoI/BamHI fragment [15] to make pEM148, and N-BfpE was expressed and purified as described for N-BfpC.

### ATPase activity

ATPase activity was measured using previously published methods [12, 16]. In brief, protein samples were added to assay buffer [20 mM Tris–HCl, pH7.6, 100 mM NaCl, 1 mM MgCl_2_, and 1 mM ATP] in a final volume of 15 μL and incubated at 37°C. Subsequently 270 μL of a mixture of 0.045% malachite green hydrochloride and 4.2% ammonium molybdate in 4 N HCl (MG/AM) was added to each sample to react with the released phosphate. After 1 min, 30 μL of citrate solution (34%) was added and mixed. The absorbance at OD_655_ was measured on a Clariostar Monochromator Microplate Reader. The ATPase activity was extrapolated from a standard curve with defined phosphate concentrations. To measure K_m_ and V_max_, various ATP concentrations were used, and the reaction was terminated at multiple time points to identify a linear data range. Data were fit by nonlinear regression to the Michaelis-Menten equation with GraphPad Prism to calculate K_m_ and V_max_. The specific activity of BfpD was measured by varying BfpD concentrations in assay buffer (above) with 2 mM ATP. The ATPase activities of N-BfpC and N-BfpE were also estimated using the same methods. To measure ATPase activities in samples of N-BfpC after gel filtration, an equal volume (8 μL in 20 μL reaction) of each fraction was tested for a period of 6 hours using 2 mM ATP.

### Site directed mutation of putative BfpD active site residues

FastCloning [17] was used to induce substitutions for glutamate codons at amino acid positions 295 and 338 of BfpD in pJZM005. In short, a pair of primers (S2Table) was designed such that each has a complementary sequence including the mutated codon and different overlapping sequences with *bfpD*. The PCR products were digested with DpnI, and subsequently transformed into *E. coli* DH5α competent cells. The desired *bfpD* mutations were confirmed by sequencing, and expressed in *E. coli* BL21(DE3) as described above.

To make plasmids for *in vivo* complementation, we first constructed a plasmid harboring wild type *bfpD* in low-copy-number cloning vector pWKS30 [18]. To do this, *bfpD* with its N terminal His tag and S tag, was isolated from pRPA405 [14] as an XbaI-SacI fragment, and inserted into pWKS30. The resultant plasmid pEMM1 was later discovered to lack its native stop codon, while a stop codon on the vector was noted downstream, thus adding an elongated non-native C-terminus to the predicted protein. We used FastCloning to restore a TAG stop codon in its original position in pEMM1, and the new plasmid was named pJZM031. Thereafter, pJZM031 was used as template to introduce E295C and E338Q mutations to obtain pJZM032 and pJZM036, respectively. FastCloning was used to introduce the E295C mutation. For the E338Q mutation, we first introduced the mutation using overlap extension PCR [19], which was later cloned into pJZM031 (XbaI/SacI digested).

### Site directed mutation of a putative BfpF active site residue

The putative active site glutamate codon at amino acid position 176 in *bfpF* was mutated to arginine, both in its native location and in the cloned gene in a plasmid used for complementation.

To make the E176R mutations, we first cloned wild-type *bfpF* into pBad24. In short, primers bfpFcom-Fwd and bfpFcom-Rev were used to clone a 1020-bp sequence including *bfpF* from the genome of EPEC strain E2348/69, and cloned into EcoRI/SalI digested pBad24. The resulting plasmid was further used as template to introduce the E167R mutation through QuikChange site-directed mutagenesis kit with primers bfpFE167R-Fwd and bfpFE167R-Rev. This construct was named pSYN125b.

A previously described allelic replacement strategy was used to introduce the E167R mutation in wild-type EPEC strain E2348/69 using suicide vector pCVD442 [20]. Primers bfpF-Fwd and bfpF-Rev were used to clone a 986 bp sequence including *bfpF* from the genome of EPEC strain E2348/69. The PCR product was subsequently cloned into SphI,SalI digested pCVD442. The resulting plasmid was further used as template to introduce the E167R mutation using the QuikChange site-directed mutagenesis kit with primers bfpFE167R-Fwd and bfpFE167R-Rev. The resulting plasmid pSYN76c was introduced into *E. coli* strain MFDpir in the presence of 0.3 mM diaminopimelic acid (DAP) and ampicillin. Donor MFDpir(pSYN76c) cells were mixed with E2348/69 for conjugation, and merodiploids were selected with ampicillin, selecting against donor cells in the absence of DAP. Clones that underwent a second recombination event were selected by growth on LB medium lacking sodium chloride with 5% sucrose, and further verified by PCR with primers bfpFE167R.F1 and bfpFE167R.R2 and sequencing with primer bfpFE167R.R2. The resulting strain with the expected E167R mutation was named UMD976. The identical method was used to introduce the BfpF_E167R_ mutation into *bfpD* mutant strain UMD926 to make strain VCU019.

### Auto-aggregation and disaggregation assay

Auto-aggregation and disaggregation were assessed as previously described [14]. Briefly, overnight cultures of E2348/69, UMD926 and VCU019 containing pWKS30, pJZM031, pJZM032, or pJZM036 were diluted 100-fold in Dulbecco’s modified Eagle’s medium and grown for 4 h at 37 °C before examination by phase-contrast microscopy. In the case of strains E2348/69, UMD916 and UMD976 containing pBad24, pSYN125b, or pJAL-F1, 0.2% arabinose was added to the medium at the time of inoculation to induce *bfpF* expression.

## Results and discussion

### Kinetics of BfpD ATPase activity in the absence and presence of partner proteins

We previously demonstrated that BfpD has ATPase activity [12]. To confirm and extend these findings, we varied the ATP concentration in the reaction and determined that the Vmax and Km of BfpD were 9.43 ± 2.21 μM min^−1^ and 1.12 ± 0.74 mM, respectively (Fig 1). The specific activity of BfpD using an ATP concentration of 2 mM was 3.66 ± 0.73 nmol min^−1^ mg^−1^ [21]. While there are many reports quantifying ATPase activity from T4P, T2SS, and other related systems (S3Table), there are few in which V_max_ and K_m_ were determined. Instead, most reported specific activity. Based on our literature review, the specific activity of similar proteins ranges from 0.71 (PilQ from thin conjugative pilus system in *E. coli*, [22]) to 37.5 nmol min^−1^ mg^−1^ (PilT from T4P in *Microcystis aeruginosa*, [23]). The activity of BfpD (3.66 nmol min^−1^ mg^−1^) falls in this range. We also attempted to purify a Hcp1-BfpD fusion protein, similar to the *V. cholerae* type II secretion ATPase EpsE Hcp1 fusion, which had 20-fold enhanced activity [4]. However, these attempts were complicated by the formation of inclusion bodies (data not shown).

**Fig 1.**
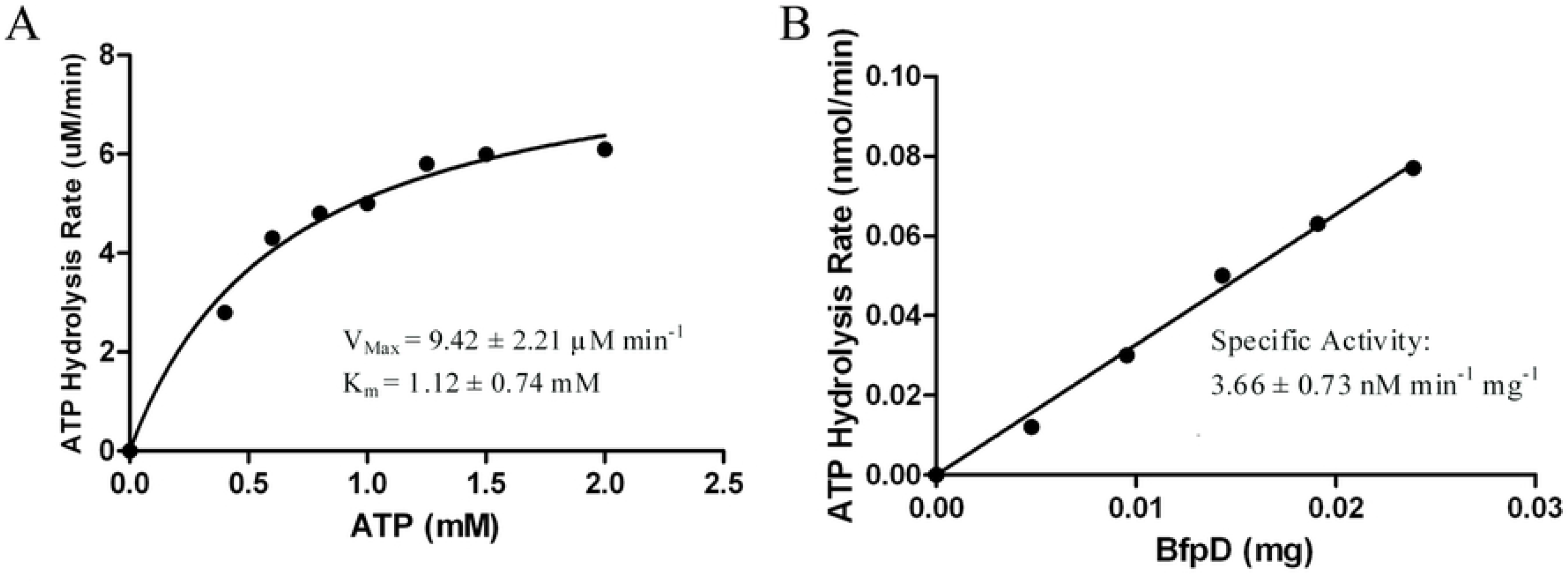
ATPase activity of purified BfpD. A) Km and Vmax: Rates of ATP hydrolysis were measured at 29.4 μM BfpD in the presence of increasing amounts of ATP. Each point represents the reaction rate derived from each ATP concentration during the linear time range. The line indicates the best fit to the Michaelis-Menten equation obtained with GraphPad Prism. B) Specific activity of BfpD was determined with increasing amounts of BfpD. The figure shown represents a typical assay, while the data shown are the mean and STDEV from 6 independent experiments.

We previously reported that BfpD activity increased approximately three-fold when co-purified with the N-terminal cytoplasmic domains of BfpC and BfpE, which together recruit BfpD to the cytoplasmic membrane [12]. To determine whether N-BfpC, N-BfpE or both proteins stimulate BfpD activity, we purified each protein separately (S1 Fig) and measured ATPase activity. Addition of N-BfpC, but not N-BfpE, to BfpD resulted in higher ATPase activity. However, we found that N-BfpC alone had ATPase activity (data not shown). This result is highly unexpected as no enzyme motifs are found in BfpC or related proteins. While PilM from *T. thermophilus* does have an ATP binding site, BfpC lacks this domain. Furthermore, PilM does not hydrolyze ATP [24]. Although the N-BfpC obtained by nickel affinity chromatography appeared to be highly pure (S1 Fig), we added an additional step to investigate whether the ATPase activity we observed was due to contamination with another protein. N-BfpC (purified from Ni-NTA resin) was applied to a HiPrep 16/60 Sephacryl S-100 column and equal volumes of each fraction were tested for ATPase activity. We found that the N-BfpC peak fractions have negligible ATPase activity, while the early fractions lacking visible protein bands have strong ATPase activity (Fig 2). This result indicates that the observed increase in BfpD ATPase activity upon addition of N-BfpC was due to contamination of N-BfpC with another ATPase.

**Fig2.**
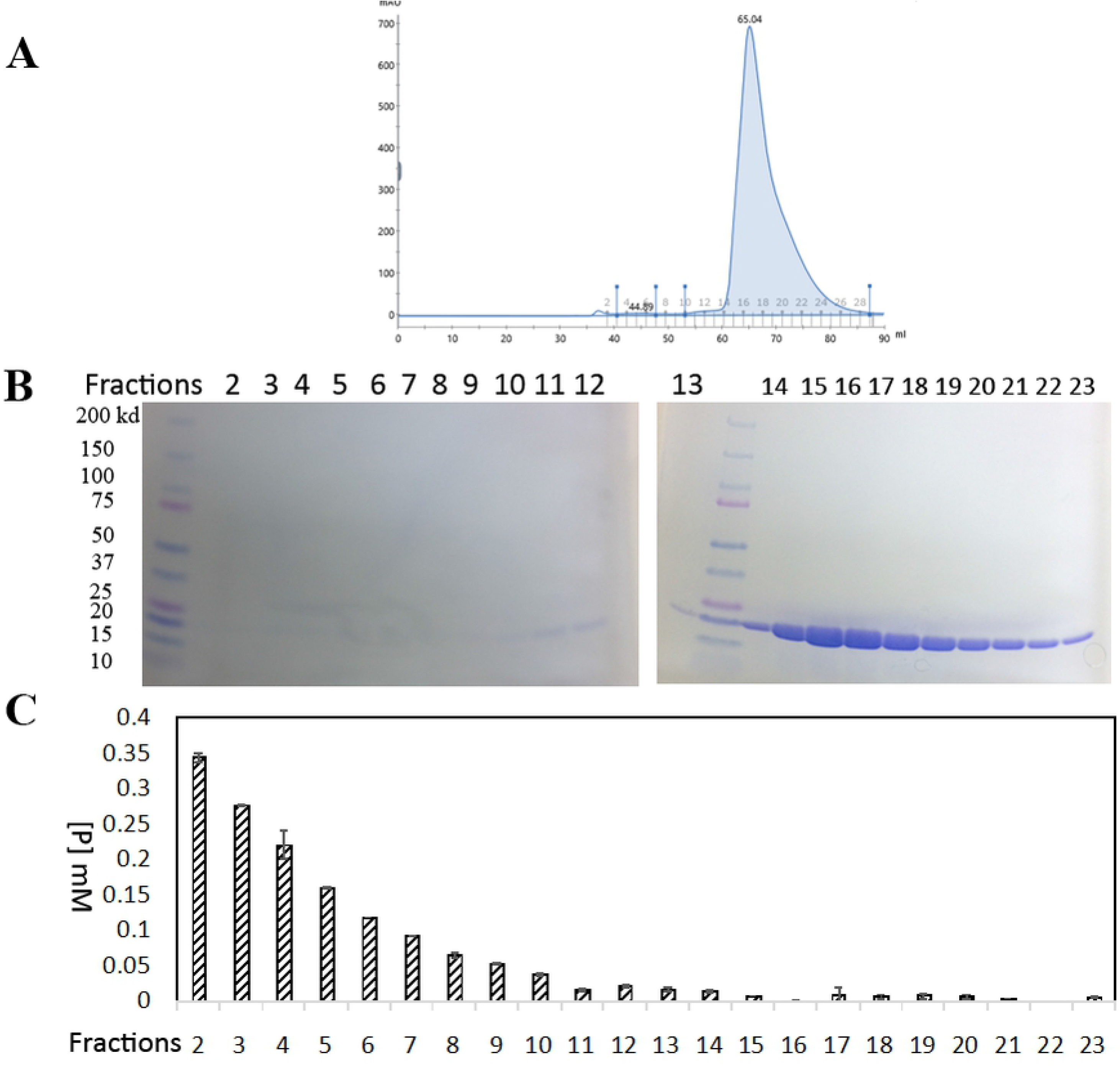
ATPase activity of N-BfpC gel filtration fractions. (A) Gel filtration elution profile of N-BfpC after nickel affinity purification (S1 Fig). (B) SDS-PAGE of 10 μL from each fraction from gel filtration. (C) ATPase activity for each fraction. For each 20 μL reaction, 8 μL of each fraction was tested for phosphate formation during a period of 6 hours using 2 mM ATP. Each test was performed three times.

### Purified BfpD variants with mutations at putative catalytic residuals retain ATPase activity

Since we determined that the observed ATPase activity of N-BfpC was due to a contaminating protein, we questioned whether the observed ATPase activity of BfpD was also from contamination, despite its apparent purity (S1 Fig). As BfpD is a member of the P-type ATPase family of large hexameric enzymes, purifying BfpD from a potential contaminating ATPase(s) through gel filtration was not feasible. We therefore mutated conserved catalytic sites in BfpD, purified these variants, and measured their ATPase activity. We chose two conserved functional residues to change (Fig 3). E295 is an absolutely conserved residue located in the Q-loop, and the counterpart in a homolog PilB (E681) from *T. thermophilus* [25] was predicted to polarize the water molecule to facilitate the nucleophilic attack on the ATP. Besides E295, E338 at the end of the Walker-B motif is also an absolutely conserved residue, and was also suggested to be involved in the hydrolysis of γ-ATP, for example, in bacterial ATPases HisP from *Salmonella typhimurium*, KpsT from *E. coli*, and MJ0796 and MJ1267 from *Methanococcus jannaschii* [26–31]). We changed E295 and E338 to several amino acids with properties similar to glutamate to minimize the chance of altering the overall structure of the protein, but avoided aspartate to reduce the possibility that the mutated protein would retain activity. Of these, we were able to express and purify BfpD_E295C_ and BfpD_E338Q_ using the protocol for native BfpD and these variants were similar in solubility. When we measured the ATPase activity of the two mutant BfpD proteins, we were able to detect ATP hydrolysis, although the V_max_ was significantly lower for both (S4Table). These results indicate either that the BfpD variants with mutations in putative catalytic residues retain some activity, or that they, and by extension native BfpD, are also contaminated with another ATPase.

**Fig 3.**
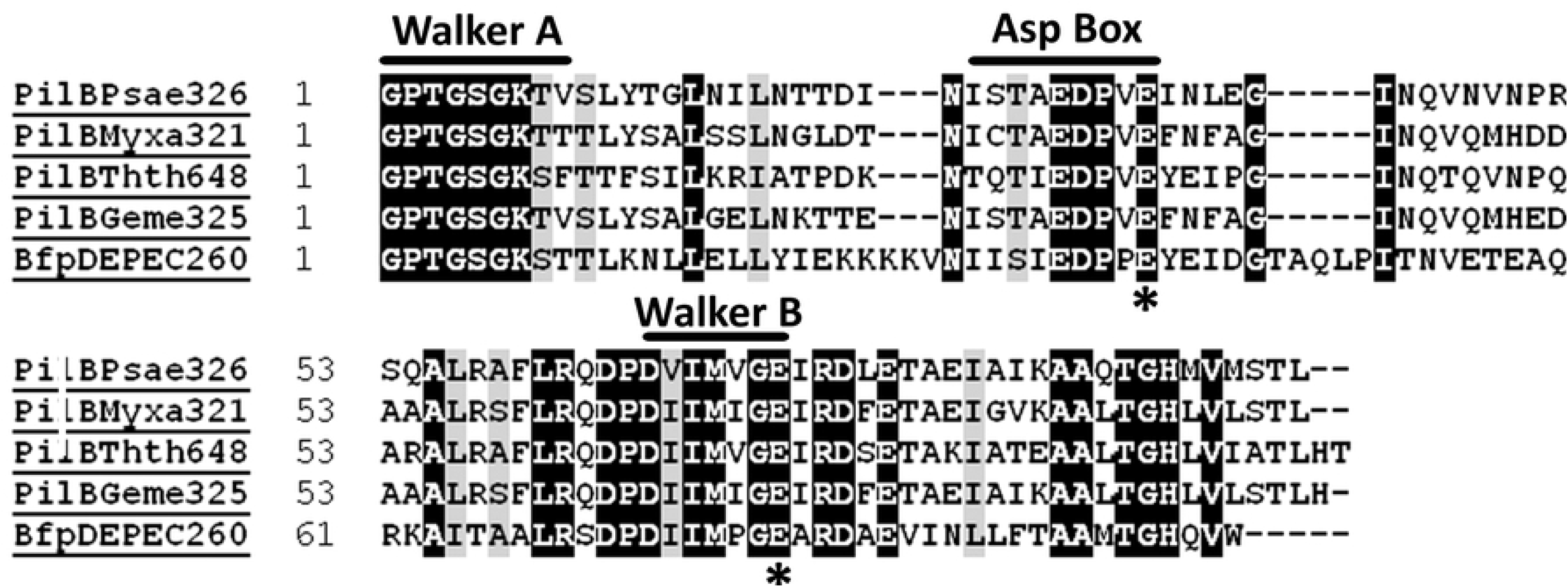
Partial amino acid sequence alignment of BfpD (BfpDEPEC260) from *E. coli* EPEC B171 with PilB homologs from *Pseudomonas aeruginosa* PAO581 (PilBPsae326), *Myxococcus xanthus* (PilBMyxa321), *Thermus thermophilus* HB8 (PilBThth648), and *Geobacter metallireducens* (PilBGeme325). Identical residues are highlighted in black, and conserved residues are highlighted in gray using a 100% shading threshold. The signature motifs are boxed and labeled. Asterisks indicate glutamates mutated in this study (E295C and E338Q). The last three numerals of the name show the starting residue position of the partial aligned sequence. For example, the BfpD partial sequence extends from the 260^th^ residue to 360^th^ residue.

### The bfpD mutants are defective in aggregation in vivo

To determine whether the E295C and E338Q *bfpD* mutants retain function *in vivo*, we evaluated their ability to complement UMD926, a *bfpD* null mutant, to restore auto-aggregation, which depends on BFP expression [32]. For this purpose, we first made pJZM031 harboring wild-type *bfpD*. From pJZM031, pJZM032 and pJZM036 were constructed, with E195C and E338Q mutations, respectively. We found that wild-type *bfpD* can restore aggregation to UMD926, as expected, while both the mutants (*bfpD*_E295C_ and *bfpD*_E338Q_) cannot (Fig 4). This experiment indicates that BfpD variants with putative catalytic mutations do not have sufficient ATPase activity *in vivo* for BFP biogenesis, further suggesting that the observed ATPase activity with the mutant BfpD proteins is, at least partly, due to contamination by (an) unknown contaminating ATPase(s). We also observed that introduction of pJZM032 and pJZM036 into wild-type strain E2348/69 had no apparent effect on the aggregation phenotype, indicating that these mutations are not dominant negative, as may have been expected if they formed inactive heterohexamers.

**Fig 4.**
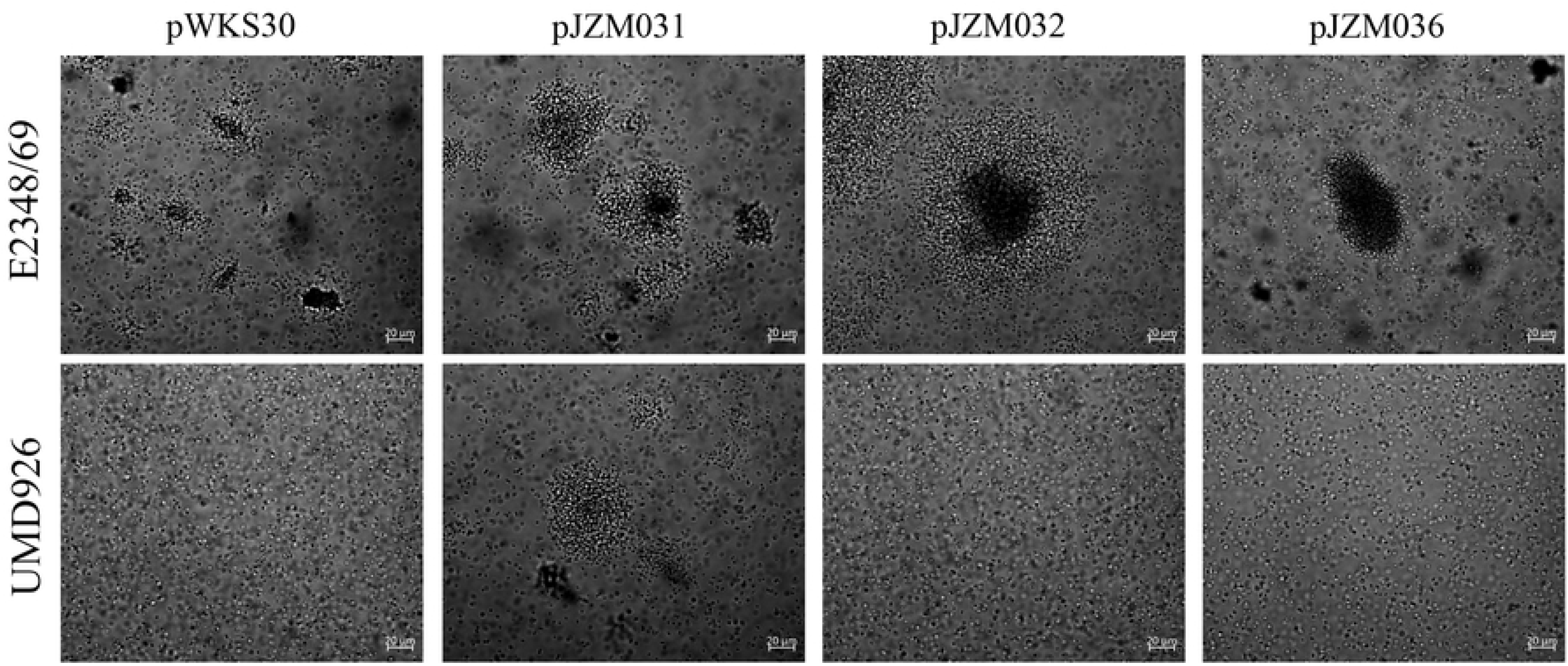
Residues E295 and E338 in BfpD are essential for function. Overnight cultures were incubated at 37° C after dilution 1:100 in DMEM. Phase-contrast micrographs were taken 4 h post-inoculation. Phase-contrast photomicrographs on the top row show wild type EPEC strain E2348/69, while those on the bottom row show *bfpD* mutant UMD926. The panels from left to right show these strains transformed with negative control pWKS30, positive control pJZM031 (wild-type *bfpD* in pWKS30), and two *bfpD* mutants pJZM032 (*bfpD_E295C_* in pWKS30) and pJZM036 (*bfpD_E338Q_* in pWKS30). Scale bars indicate that all images have the same magnification.

### BfpF_E167R_ is dominant negative

When we created a strain (UMD976) that has a putative catalytic mutation (E167R) in the gene for retraction ATPase BfpF that is analogous to the *bfpD*_E295C_ allele, we were unable to complement this mutant with wild type *bfpF* in plasmid pJAL-F1 (Fig 5). In contrast, the results above show that active site mutations of BfpD (E195C and E338Q) do not interfere the function of wild-type BfpD. To determine whether the *bfpF_E195R_* mutation is dominant negative, we constructed plasmid pSYN125b, which harbors this mutation. Wild-type strain E2348/69 failed to disaggregate when transformed with pSYN125b (Fig 5), indicating that BfpF_E167R_ interferes with the normal function of wild-type BfpF. In contrast, pJAL-F1 can restore the normal disaggregation ability of UMD916, in which the *bfpF* gene is disrupted by insertion of a non-polar antimicrobial resistance cassette [33] (Fig 5). These results indicate that catalytic mutations in *bfpF*, in contrast to similar mutations in *bfpD*, are dominant negative.

**Fig 5.**
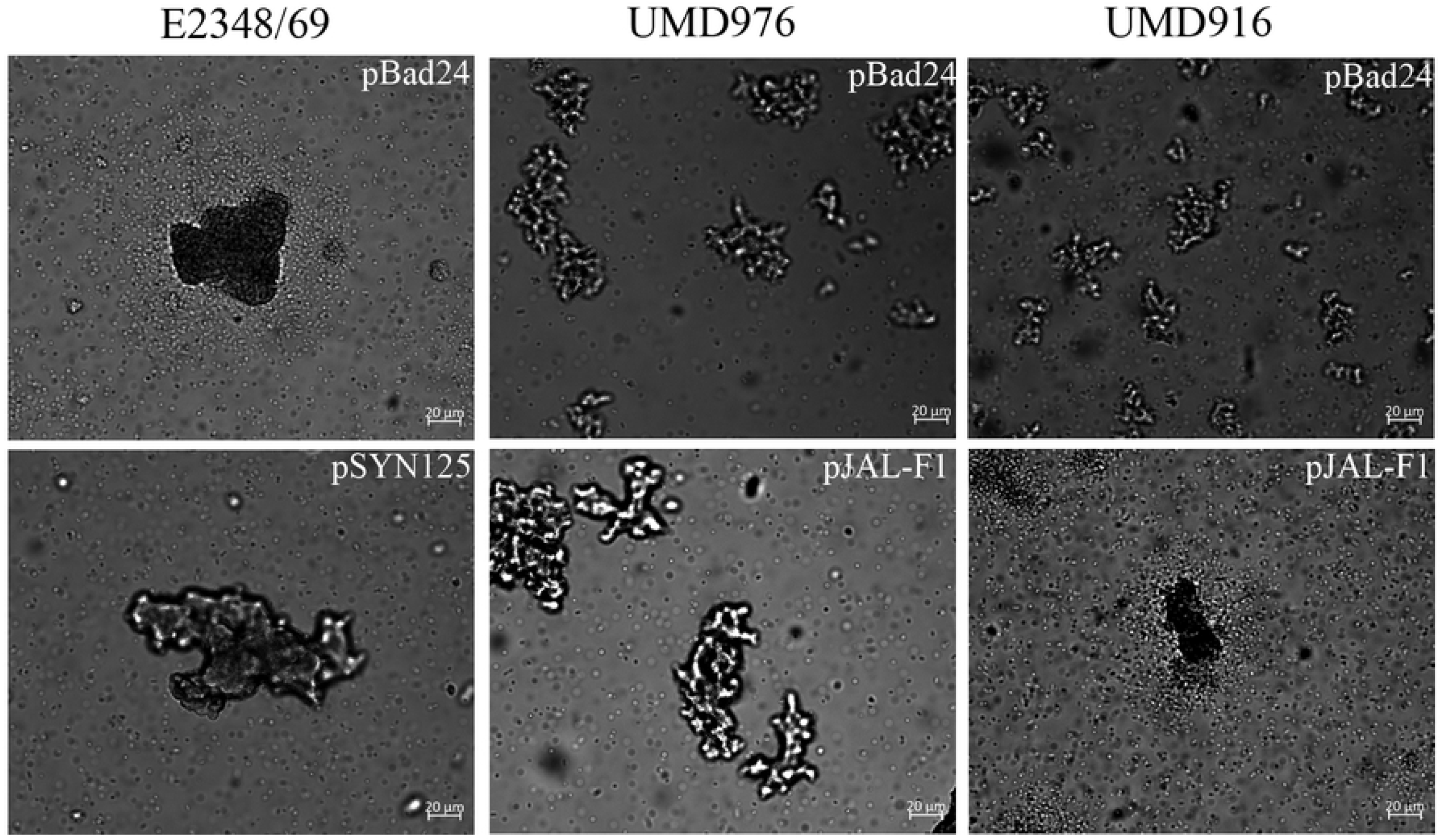
Residue E167 in BfpF is essential for function, and E167R causes a dominant negative effect on disaggregation. Overnight cultures were incubated at 37° C after dilution 1:100 in DMEM plus 0.2% arabinose for 4 h. Strain names are shown on top: E2348/69 (wild-type EPEC strain), UMD976 (E2348/69 *bfpF*_E167R_ site mutation), and UMD916 (*bfpF::aphA3* null mutant). Strains in the top panel were transformed with control vector pBad24, while those in the bottom row were transformed with pBad24 containing *bfpF*_E167R_ (pSYN125b) or wild-type *bfpF* (pJAL-F1). While all panels show auto-aggregation, only E2348/69 (pBad24) and UMD 916 (pJAL-F1) demonstrate individual bacteria surrounding aggregates, indicative of dis-aggregation. Scale bars indicate that all images have the same magnification.

### The *bfpD* catalytic mutants remain defective in aggregation in the absence of a retraction ATPase

Although *bfpD* mutations at E195 and E338 cannot restore aggregation to a *bfpD* null mutant, it remains possible that these BfpD variants still possess some residual activity, since markedly reduced activity may be undetectable in the presence of the retraction forces exerted by BfpF. We therefore examined the ability of the plasmids encoding variants of BfpD to restore aggregation in the absence of BfpF. For this purpose, wild-type *bfpF* was replaced with *bfpF* encoding BfpF_E167R_ in *bfpD* mutant UMD926 to create the double *bfpD bfpF* mutant VCU019. Subsequently, plasmids pJZM031, pJZM032, pJZM036 and vector pWKS30 were transformed into VCU019. Among these strains, only VCU019 (pJZM031) encoding wild type BfpD demonstrated an aggregation phenotype without disaggregation (Fig 6). None of the other strains showed aggregation. This result further strengthens our conclusion that BfpD_E295C_ and BfpD_E338Q_ have no residual activity *in vivo*, and the residual ATPase activity observed *in vitro* must be from occult contamination by a highly active ATPase.

**Fig 6.**
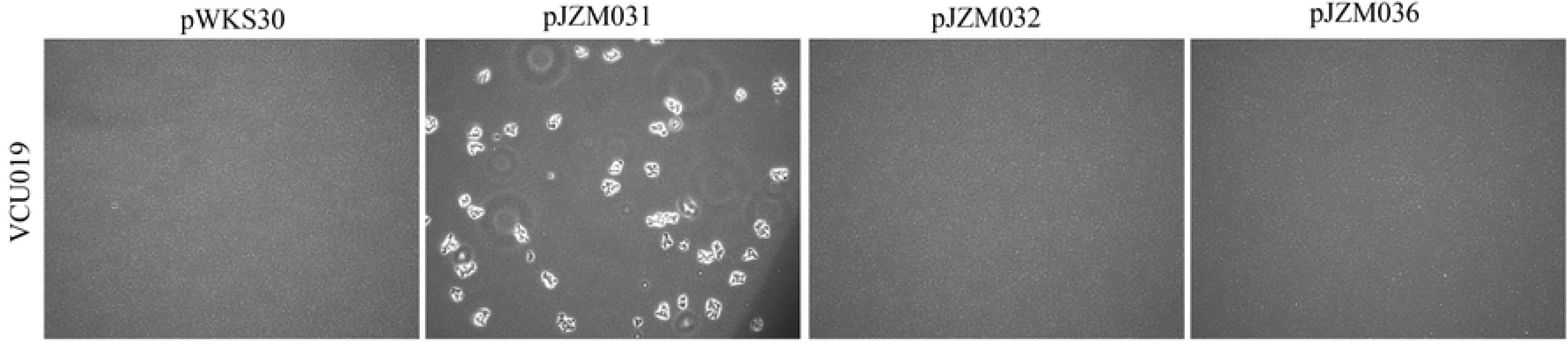
Residues E295 and E338 in BfpD have no residual function *in vivo*. Overnight cultures were incubated at 37° C after dilution 1:100 in DMEM. Phase-contrast micrographs Strain VCU019 was derived from *bfpD* mutant strain UMD926 and additionally has the *bfpF_E167R_* allele in place of wild type *bfpF*. VCU019 was transformed with negative control vector pWKS30, positive control pJZM031 containing wild-type *bfpD* in pWKS30, plasmid pJSM032 containing *bfpD_E295C_* in pWKS30, or pJZM036 containing *bfpD_E338Q_* in pWKS30, as indicated. All images have the same magnification.

### Perspectives

This study illustrates unique characteristics of the BFP T4P system, emphasizing that results from similar systems cannot be assumed to apply to all others. For example, while the addition of EspL, a structural homologue of N-BfpC, to T2SS ATPase EpsE enhanced the activity of the latter [5–6], we were unable to detect any effect of either N-BfpC or N-BfpE on the *in vitro* activity of BfpD. And while a putative catalytic mutation in the retraction ATPase BfpF has a dominant-negative phenotype, a similar mutation in extension ATPase BfpD does not. This study also delivers the cautionary message that sensitive enzymatic activity assays may belie what appears on SDS-PAGE to represent proteins purified to homogeneity. Further studies of BfpD enzyme activity will require additional purification steps.

## Acknowledgments

We are grateful to Ekatarina Milgotina (currently at GSSHealth, Baltimore, MD) and Kurt Piepenbrink (currently at University of Nebraska) for creating plasmids pEM148 and pBfpD-HCP, respectively.

## Supporting Information

**S1 Fig**. N-BfpC (1), BfpD (2), and N-BfpE (3) were separated by 4-15% SDS-PAGE in parallel with protein standards (Precision Plus Protein™ Dual Color Standards, #1610374, Biorad). The gel was stained with Coomassie Brilliant Blue R-250 dye.

**S1 Table. Strains, and plasmids used in this study.**

**S2 Table. Oligonucleotides used in this study.**

**S3 Table. Published ATPase activities of proteins related to BfpD.**

**S4 Table. V_max_ for wild-type BfpD, BfpD_E295C_, and BfpD_E338Q_.**

